# Machine-learning-guided library design cycle for directed evolution of enzymes: the effects of training data composition on sequence space exploration

**DOI:** 10.1101/2021.08.13.456323

**Authors:** Yutaka Saito, Misaki Oikawa, Takumi Sato, Hikaru Nakazawa, Tomoyuki Ito, Tomoshi Kameda, Koji Tsuda, Mitsuo Umetsu

## Abstract

Machine learning (ML) is becoming an attractive tool in mutagenesis-based protein engineering because of its ability to design a variant library containing proteins with a desired function. However, it remains unclear how ML guides directed evolution in sequence space depending on the composition of training data. Here, we present a ML-guided directed evolution study of an enzyme to investigate the effects of a known “highly positive” variant (i.e., variant known to have high enzyme activity) in training data. We performed two separate series of ML-guided directed evolution of Sortase A with and without a known highly positive variant called 5M in training data. In each series, two rounds of ML were conducted: variants predicted by the first round were experimentally evaluated, and used as additional training data for the second-round prediction. The improvements in enzyme activity were comparable between the two series, both achieving enzyme activity 2.2–2.5 times higher than 5M. Intriguingly, the sequences of the improved variants were largely different between the two series, indicating that ML guided the directed evolution to the distinct regions of sequence space depending on the presence/absence of the highly positive variant in the training data. This suggests that the sequence diversity of improved variants can be expanded not only by conventional ML using the whole training data, but also by ML using a subset of the training data even when it lacks highly positive variants. In summary, this study demonstrates the importance of regulating the composition of training data in ML-guided directed evolution.

## Introduction

Proteins are a class of biopolymer in which amino acids with various side chains are linked by peptide bonds. The diversity of possible amino acid sequences generates various biogenic functions, and allows us to create novel functions not found in nature. While genomics studies have provided vast information on natural proteins, they cover only a small subset of sequence space (defined as the set of all possible amino acid sequences of a protein). Thus, mutagenesis-based protein engineering has been used to create novel protein variants with desired functions by introducing amino acid mutations into natural proteins, which is called directed evolution [1].

Current experimental approaches for directed evolution often fail to obtain desirable variants due to the difficulty in exploring sequence space. Iterative saturation mutagenesis (ISM) is a conventional method where saturation mutagenesis (substitution of one or a few residues for all possible amino acids) and selection of the optimal variant are iteratively conducted [2]. In this method, the explored region of sequence space at each iteration may be too small to contain desirable variants with high probability. This probability can be increased by preparing an extremely large variant library with mutations in many residues while the library size would be beyond the limit of comprehensive screening [3-5]. Therefore, the success of directed evolution crucially depends on preparing a small library with high enrichment of desirable variants.

Recently, machine learning (ML) has been used to accelerate directed evolution of proteins [6,7]. In this method, saturation mutagenesis and/or random mutagenesis are performed to generate an initial library. The variants in the library are experimentally evaluated to obtain their sequences and functions, and then used as training data to construct a ML model that predicts the function from the sequence. By using the ML model, a second-round library that contains variants predicted to have improved functions is generated. This method enables to design a library with high enrichment of desirable variants, and thus has been successfully applied to directed evolution of various proteins including fluorescent proteins [8-10], enzymes [11-13], and others [14,15]. However, it remains unclear how ML-guided directed evolution explores sequence space depending on the composition of training data. Will different training data lead to different regions of sequence space, and different levels of functional improvements?

Here, we perform ML-guided directed evolution with different training data to investigate the differences in explored regions of sequence space and resultant functional improvements. As an example, we used Sortase A (SrtA), a transpeptidase enzyme found in Gram-positive bacteria. SrtA conjugates a peptide containing an LPXTG sequence to another peptide containing a GGG sequence [16]. This sequence-specific transpeptidase activity has been utilized for protein modification, and conjugation of proteins to proteins, polymers, and solid substrates in *in vivo* and *in vitro* [17-21]. Although the wild-type SrtA has poor reaction kinetics that limit its application, several variants with superior activity have been reported [18,22,23]. To make different training data for ML, two typical scenarios encountered in protein engineering were considered: (1) the situation where we already have a “highly positive” variant (variant with a desired function) at least to some degree; and (2) the situation where no such variant is available. Accordingly, we conducted two separate series of ML-guided directed evolution with and without a highly positive SrtA variant called 5M [22] in training data. Interestingly, variants with higher enzyme activity than 5M were discovered by both series with comparable improvements (2.2–2.5 times higher than 5M) while the sequences of the improved variants were largely different between the two series. This demonstrates the ability of ML to guide directed evolution to distinct regions of sequence space depending on the composition of training data, which is useful to expand the sequence diversity of improved variants.

## Results

### ML-guided library design cycle

In our previous study [8], we proposed a ML-guided directed evolution method for fluorescent proteins. In this method, several residues at certain positions in a target fluorescent protein were mutated to make a variant library for training a Bayesian ML model, and then the ML model was used to rank all possible variants according to their predicted probability of having improved fluorescence performance. Here, we extend this method to enzymes with iterative design of libraries (Figure 1; Methods). In brief, certain residues in a target enzyme are mutated to make an initial library, and functional data (sequence, expression level, and enzyme activity) for approximately 100 variants in this library are obtained. These data are then used to train a ML model, and top-ranked variants predicted by ML are used to design the second-round library. The functional data for dozens of variants in the second-round library are obtained, and used as additional training data for the ML model to design the third-round library. We applied this ML-guided library design cycle to SrtA for improving enzyme activity.

**Figure 1.**
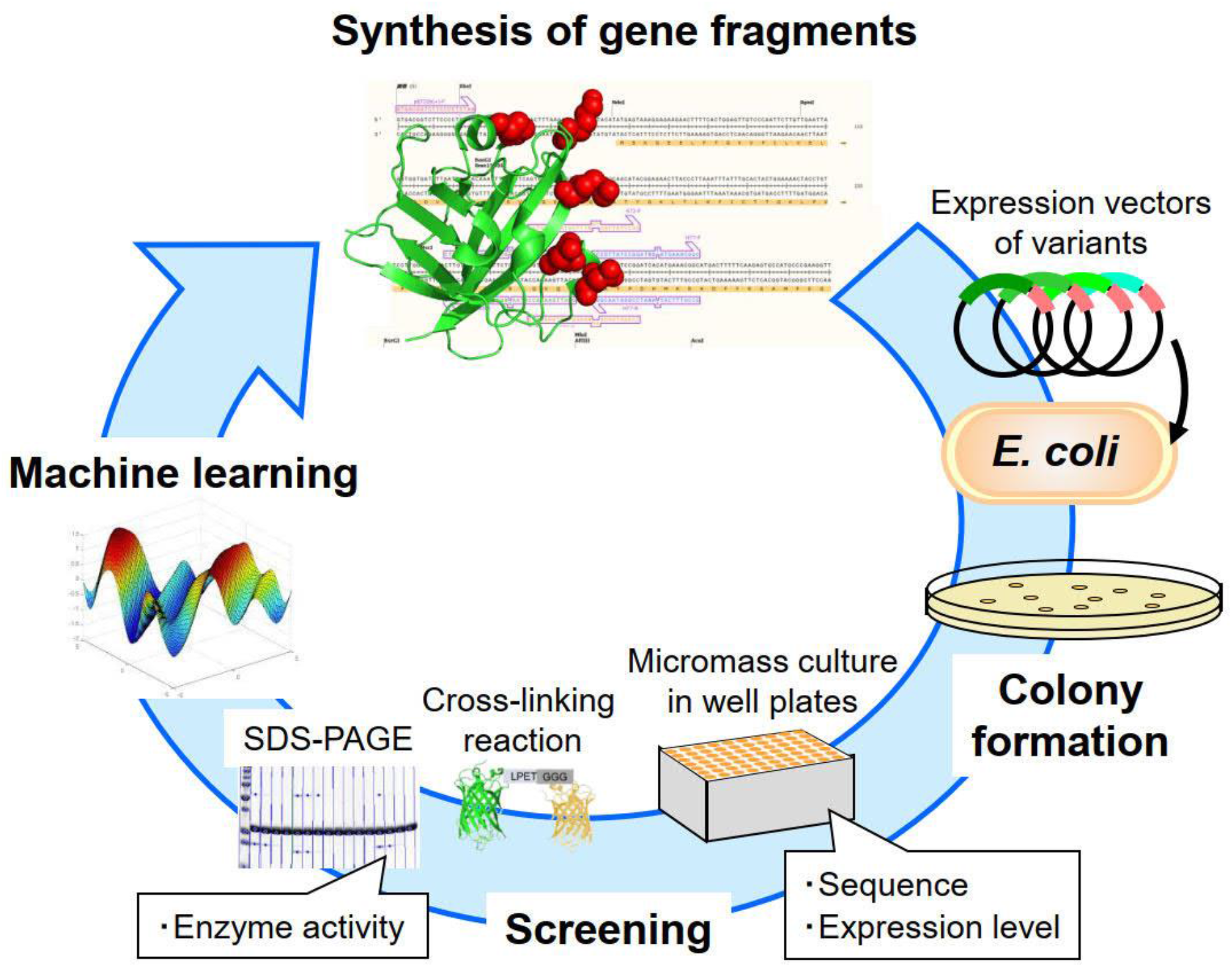
Overview of ML-guided library design cycle for directed evolution of SrtA. See the main text for the detail explanation.

### Preparation of the initial library

To select the residues that should be mutated in SrtA, we referred to the reported high activity variant called 5M [22] that has five mutated residues: P94R, D160N, D165A, K190E, and K196T (Figure 2). At these five residues, point mutagenesis and site-directed random mutagenesis were applied to make the initial library. The expression vectors for this library were prepared as described in our previous study [8], and used to transform *Escherichia coli*. Transformants were cultured in a 96-deep-well plate, and the sequence of each variant was determined. Then, each variant was purified by immobilized metal affinity chromatography (IMAC), and the expression level was measured. Variants with sufficient expression levels were used for enzyme activity assay. In this assay, two substrates—cycle3 green fluorescent protein (GFP) with an LPETG sequence at the C-terminus and Venus yellow fluorescent protein (YFP) with a GGG sequence at the N-terminus—were conjugated with each other by a SrtA variant [18] (Supplementary Figure S1), and the enzyme activity was estimated by SDS-PAGE analysis. In total, 80 variants were prepared (56 from site-directed random mutagenesis and 24 from point mutagenesis). The enzyme activity of 45 of these variants was measured while the expression levels of the 35 other variants were insufficient for measuring enzyme activity. The sequences of these variants are shown in Supplementary Table S1. Most variants had low expression levels and enzyme activity (Figure 3); only four of the 45 measured variants showed higher enzyme activity than the wild type, and all variants showed lower enzyme activity than 5M. These data were used to train a ML model.

**Figure 2.**
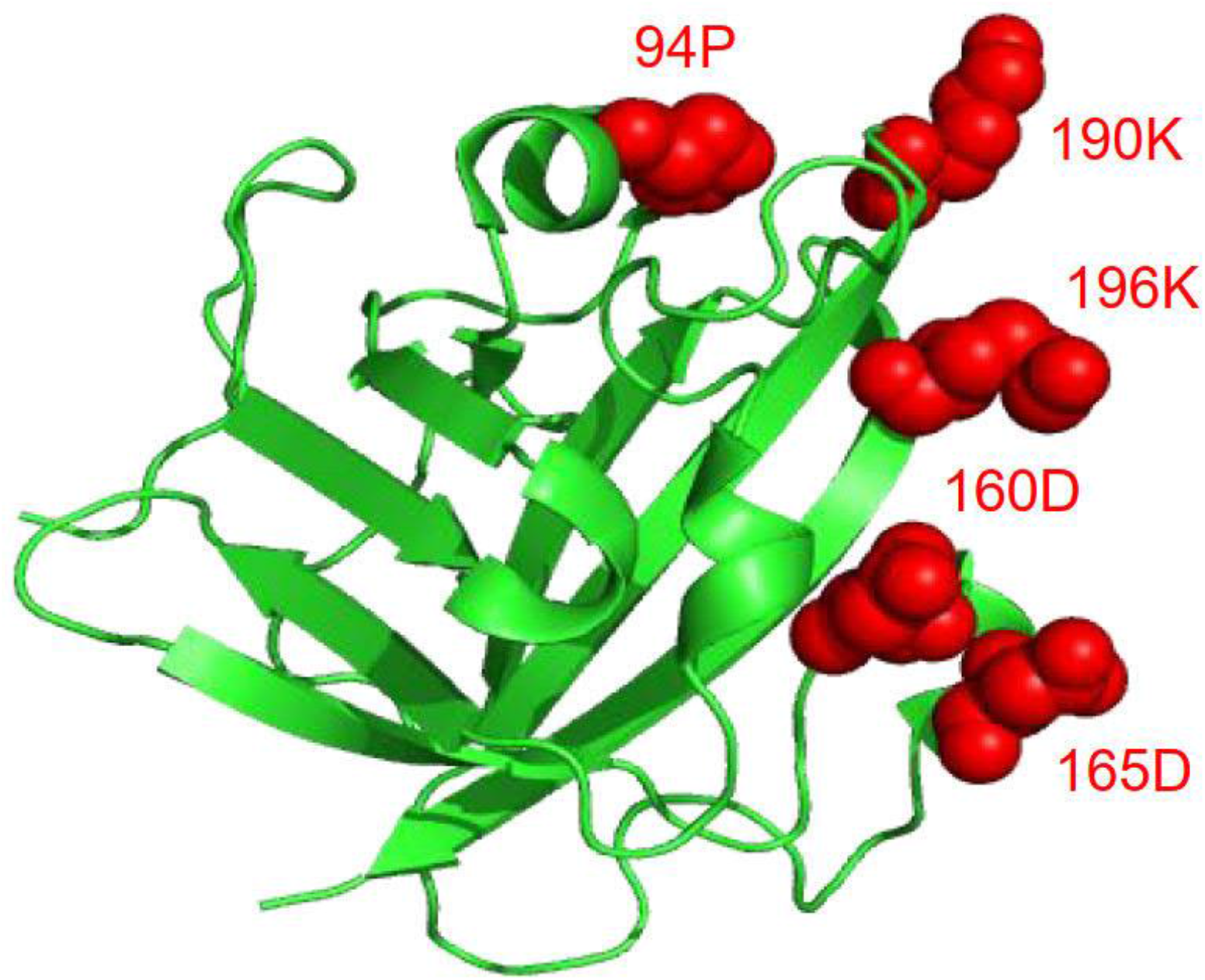
Structure of SrtA (PDB ID: 1T2P). The five mutated residues (94P, 160D, 165D, 190K, and 196K) are colored in red.

**Figure 3.**
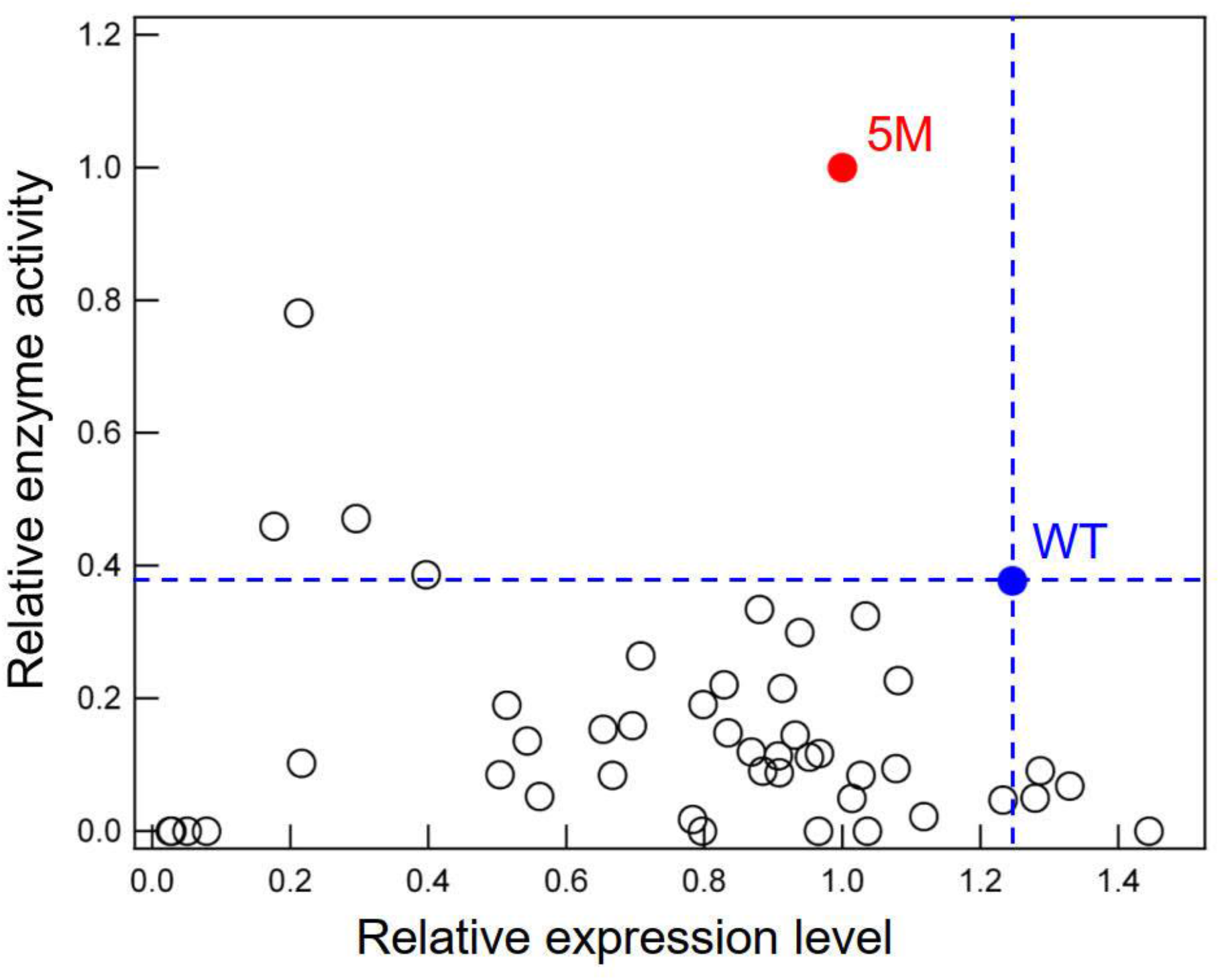
Expression levels and enzyme activity of the SrtA variants in the initial library. Values are normalized by those of 5M. Dashed lines represent the values of the wild type (WT). Only variants with measured enzyme activity (Supplementary Table S1) are shown.

### ML with the initial library

We constructed a ML model based on a Gaussian process that predicts the enzyme performance score of a SrtA variant from its sequence (Methods; Equations 1 and 2). The enzyme performance score is a sigmoidal function of enzyme activity while it takes zero when the expression level is insufficient for measuring enzyme activity (e.g., the 35 low-expressed variants in Supplementary Table S1). This definition of the enzyme performance score allows the ML model to predict variants with high enzyme activity while avoiding variants with insufficient expression levels. For feature vectors used in the ML model, we considered a variety of amino acid descriptors based on physicochemical properties, structural topology, and evolutionary information as well as their combinations. In benchmark experiments, we found that the combination of Z-scale [24] and position-specific score matrix (PSSM) [25] achieved the best prediction accuracy (Methods; Supplementary Figure S4; Supplementary Table S4). The dimensionality of this feature vector was six per residue. Thus, the number of features used in our final model was 30 (i.e., 6 dimensions × 5 mutated residues).

To train the ML model, two different datasets were prepared (Table 1): one consisted of the 80 variants from the initial library together with the wild type and 5M (initial 5M+ library); and the other consisted of the 80 variants and the wild type but excluded 5M (initial 5M− library). These datasets reflected practical scenarios in protein engineering: the 5M+ dataset corresponds to the situation where we already have a highly positive variant for a target protein (e.g., 5M for SrtA), and hope to discover superior variants; and the 5M− dataset simulates the situation where no such variant is available. We constructed two ML models by using the 5M+ and 5M− datasets, respectively, as training data. By using each ML model, we ranked all possible variants regarding the five mutated resides (i.e., 20^5^ = 3,200,000 variants) excluding those in the training data (Supplementary Data S1 and S2 for 5M+ and 5M−, respectively). This experimental design allowed us to evaluate the effects of a highly positive variant on prediction results in ML-guided directed evolution.

**Table 1.** Summary of the initial library. The numbers of SrtA variants used as training data for ML are shown. The initial 5M+ library contains a highly positive variant 5M while the initial 5M− library does not. Activity measured: variants with measured enzyme activity. Low expression: variants with insufficient expression levels for measuring enzyme activity. See Supplementary Table S1 for the sequence information.

|  | Initial 5M+ library | Initial 5M- library |
| --- | --- | --- |
| Activity measured | 45 | 45 |
| Low expression | 35 | 35 |
| Wild type | 1 | 1 |
| 5M | 1 | 0 |
| Total | 82 | 81 |

In the prediction result for 5M+, the six high activity variants reported previously [22] were ranked within the top 2% (Table 2), implying that the ML prediction was reliable. The prediction result for 5M− showed comparable reliability except that 5M, which has been reported to have the highest activity among these six variants [22], had a slightly low rank within the top 5%.

**Table 2.**
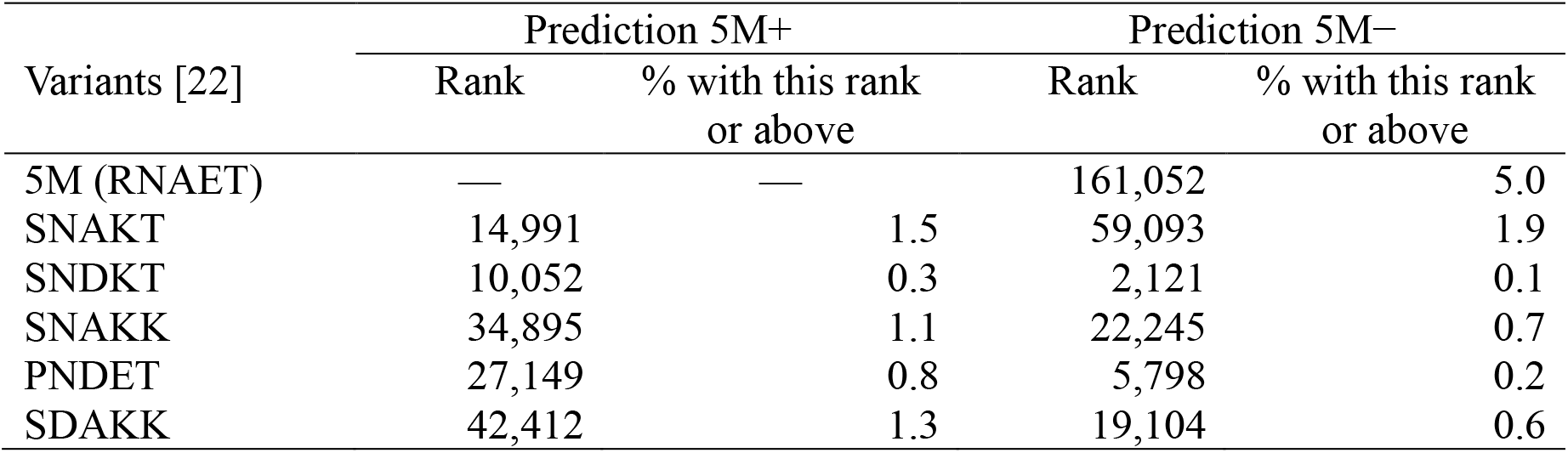
Ranks of the reported high activity SrtA variants by ML prediction. Ranks among all possible variants (20^5^ = 3,200,000) excluding those in the training data are shown. “Prediction 5M+” and “Prediction 5M−” represent the prediction results by the ML models trained with the initial 5M+ and 5M− libraries, respectively. Variants are represented by amino acids at the five mutated residues (94, 160, 165, 190, and 196).

### Screening the variants in the second-round library

To assess the performance of the predicted variants, the second-round library was made for each of 5M+ and 5M− by means of the mix primer method (Methods). The primer sequences were designed so that the top 50 variants in the ranking list were contained in the library. After cloning, we could prepare 37 variants ranked within the top 50 plus 30 variants ranked within 51–606 for the second-round 5M+ library, and 45 variants within the top 50 plus 31 variants ranked within 52– 4238 for the second-round 5M− library (Supplementary Table S2). Interestingly, most of these variants (except that ranked 29th in the ranking list of 5M−) were prepared with sufficient expression levels for measuring enzyme activity. This suggests that the enzyme performance score used in the ML model, which we designed to incorporate not only enzyme activity but also expression level (Methods; Equation 2), contributed to the prediction of variants with superior expression levels compared to those in the initial library.

For both second-round 5M+ and 5M− libraries, many variants showed higher enzyme activity than the wild type (Figure 4). Three variants in the second-round 5M− library showed higher enzyme activity than 5M while a few variants in the second-round 5M+ library showed comparable enzyme activity to 5M (Supplementary Figure S2). These results indicate that ML successfully designed the second-round libraries with high enrichment of high activity variants.

**Figure 4.**
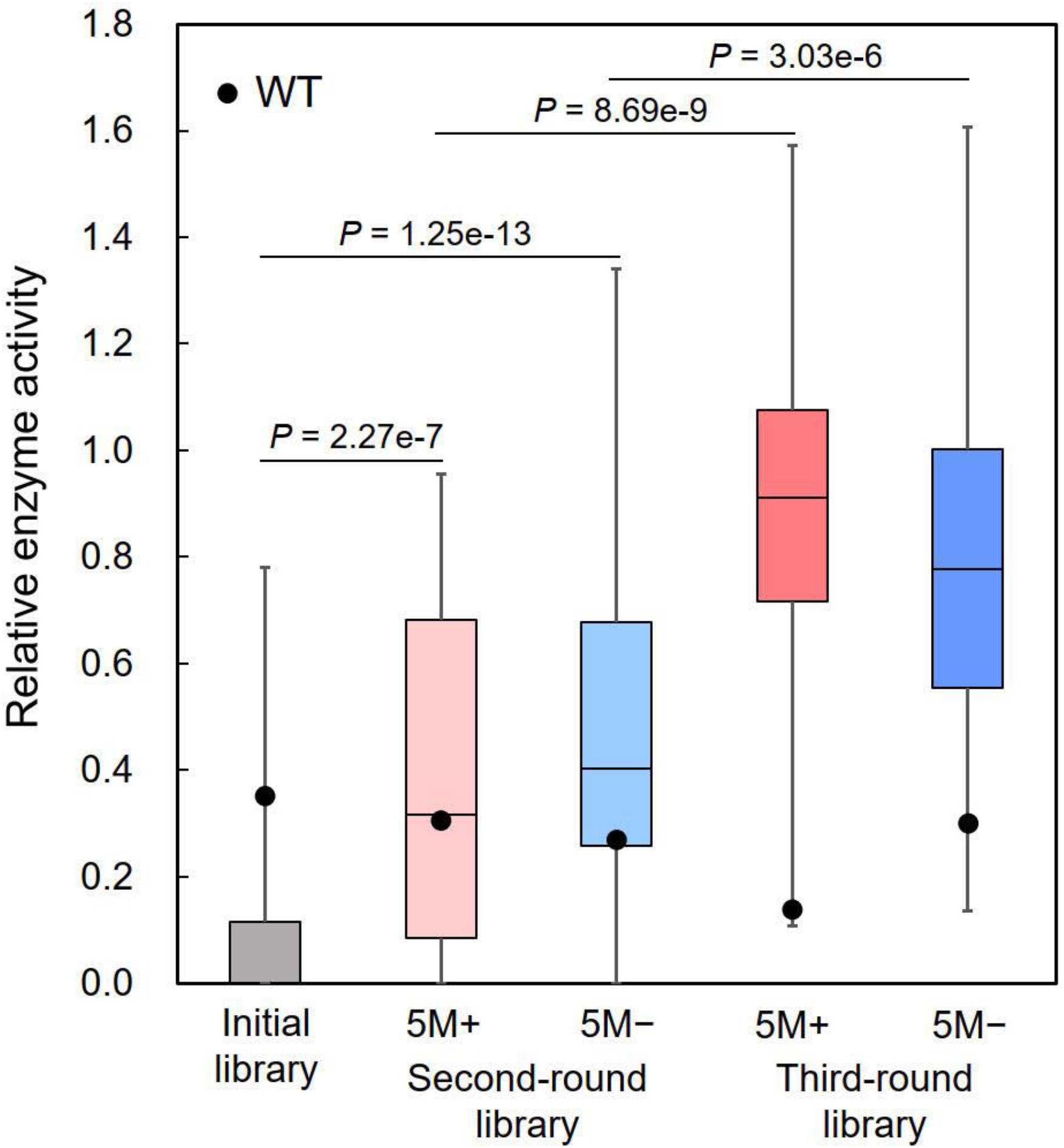
Enzyme activity of the SrtA variants in the initial and second- and third-round libraries. Values are normalized by that of 5M. Black dots represent the values of the wild type (WT). In all box plots, 5M and WT are not included in their distributions. Variants with insufficient expression levels for measuring enzyme activity are plotted as enzyme activity of zero; in the box plots of the initial and second-round libraries, lower bounds are zero due to these variants. *P*-values were calculated by one-sided Mann-Whitney *U* test.

### Iteration of ML-guided library design

The results of the second-round libraries were used as additional training data for the ML models, and the third-round libraries were prepared by the same method as used for the second-round libraries (Supplementary Data S3-S4; Supplementary Table S3). Among these, only four variants showed insufficient expression levels for measuring enzyme activity (Supplementary Table S3). Most variants showed higher enzyme activity than the wild type (Figure 4), of which 19 and 12 variants in the case of 5M+ and 5M−, respectively, were also higher than 5M (Supplementary Figure S3). Thus, the iteration of ML-guided library design dramatically increased the fraction of high activity variants, enabling the discovery of many variants with superior activity to 5M.

We evaluated the enzyme activity of the improved variants in detail. For both 5M+ and 5M− libraries, we selected the top variant from the second-round libraries and the top three variants from the third-round libraries in terms of enzyme activity measured in the above screening process. These variants were individually prepared by culture in 1-L flasks, and purified by IMAC and size exclusion chromatography (SEC). The enzyme activity of the purified variants was measured under various concentrations of the substrate GFP and a single concentration of the substrate YFP (Figure 5); the range of GFP concentrations extended higher than that used in the above screening process. All the variants showed higher enzyme activity than the wild type and 5M. In particular, the variants from the third-round libraries showed the enzyme activity 4.4–5.0 and 2.2–2.5 times higher than the wild type and 5M, respectively, at the highest GFP concentration tested (120 µM).

**Figure 5.**
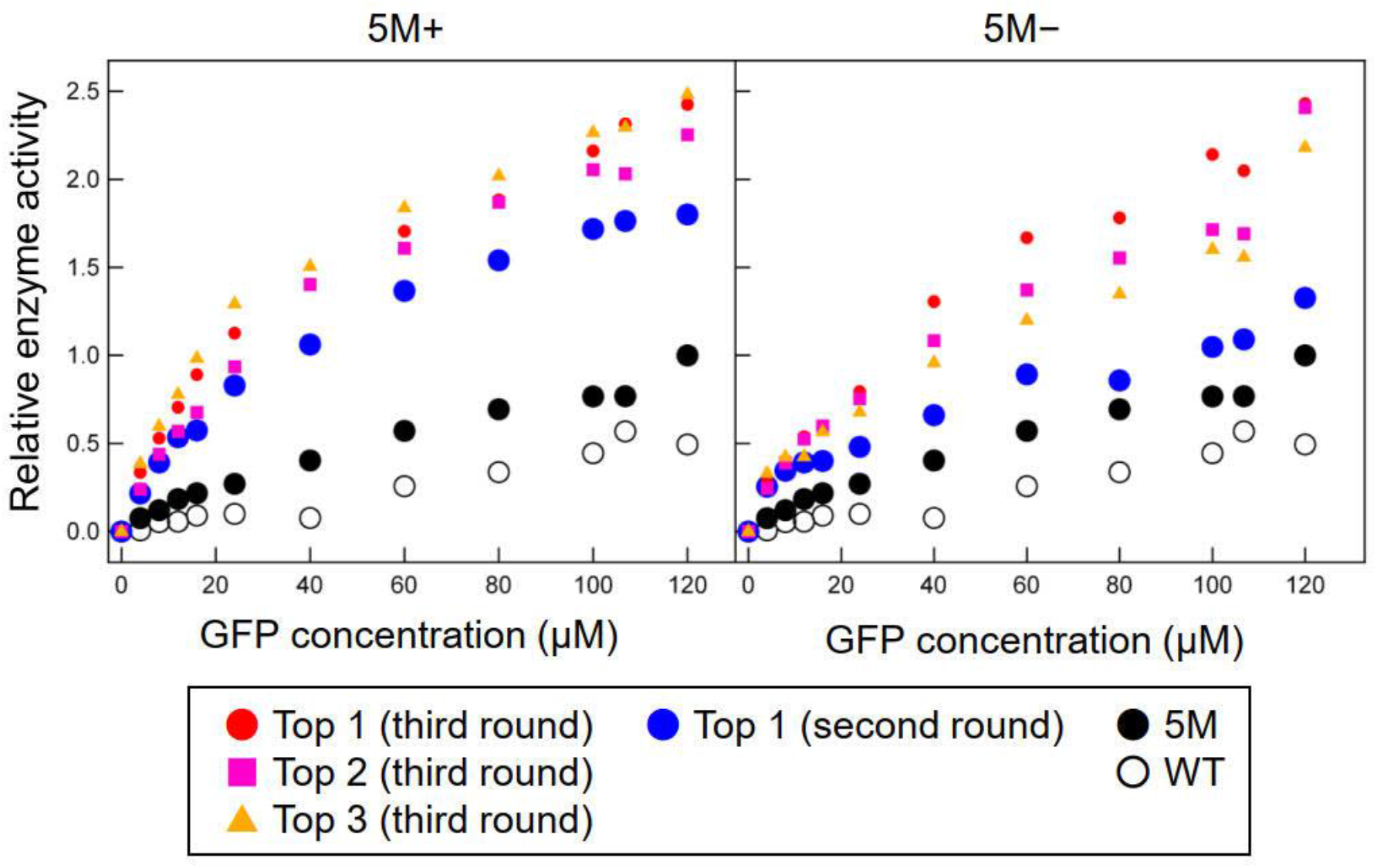
Enzyme activity of the SrtA variants evaluated under different concentrations of the substrate GFP. The concentration of the substrate YFP is fixed at 200 μM. Enzyme activity is normalized by that of 5M at the highest GFP concentration (120 μM). The top variant from the second-round libraries and the top three variants from the third-round libraries (in terms of measured enzyme activity) are shown. WT: wild type.

Remarkably, the improvements in enzyme activity were comparable between the 5M+ and 5M− libraries. This suggests that ML-guided directed evolution can discover improved variants even when training data lack highly positive variants such as 5M for SrtA.

### Trajectory of directed evolution in sequence space

Despite the comparable improvements in enzyme activity, the sequence profiles of the variants were largely different between the 5M+ and 5M− libraries (Figure 6A). In the case of 5M+, the amino acids appearing in the 5M sequence were frequently found in the variants from the second-round library, but their frequencies tended to become lower in the third-round library. Concomitantly, in the case of 5M−, the amino acids in the variants from the second-round library were dominated by those appearing in the wild type sequence, and the frequencies of these amino acids tended to be lower in the third-round library.

**Figure 6.**
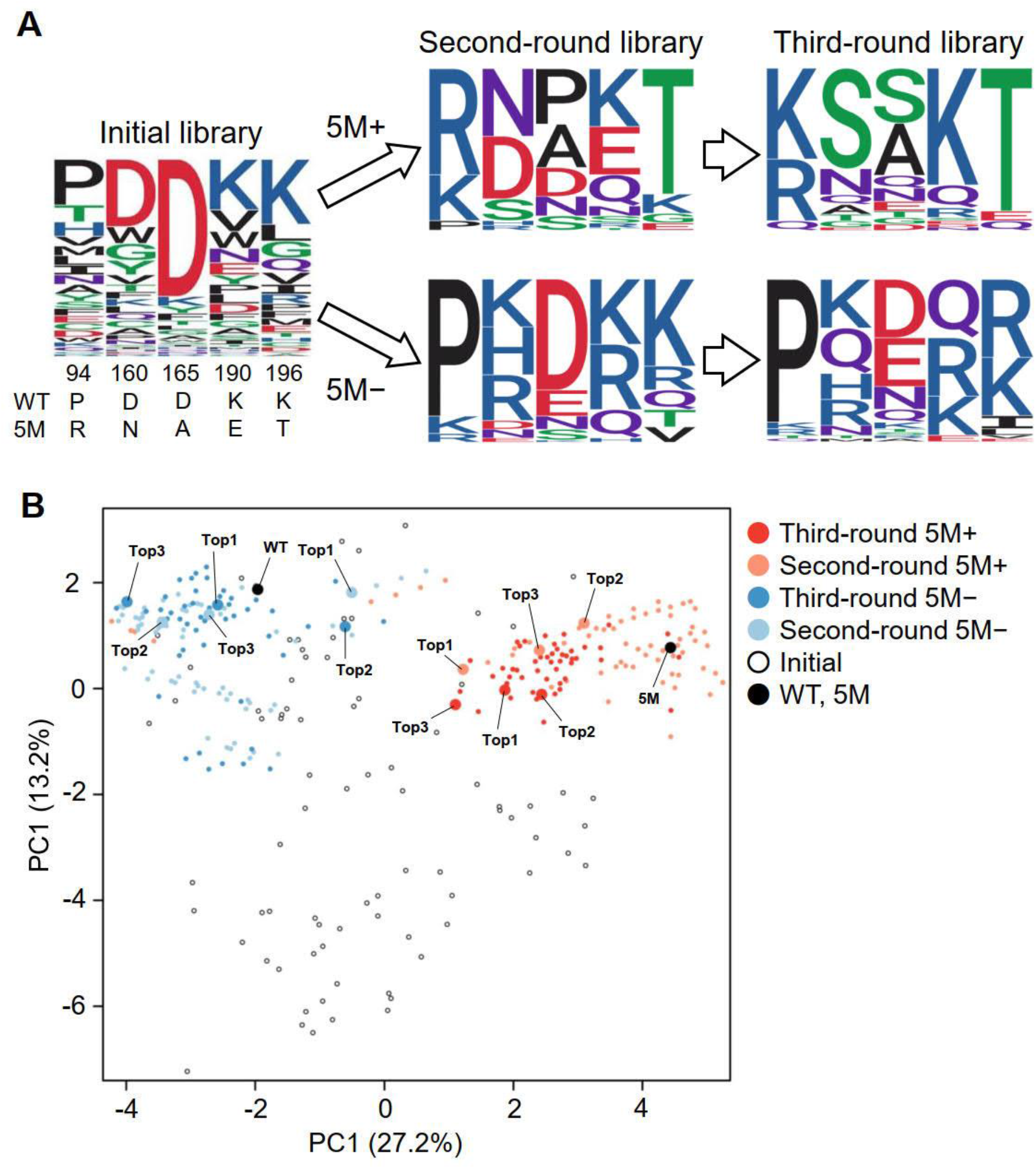
Visualization of the ML-guided directed evolution of SrtA in sequence space. (A) Sequence logo representation of amino acids at the five mutated residues in the initial and second- and third-round libraries. The amino acids in the wild type (WT) and 5M are shown below the sequence logo of the initial library. (B) Principal component analysis of the variants in the initial and second- and third-round libraries. WT, 5M, and the top 3 variants (Top1, Top2, Top3) in the second- and third-round libraries (in terms of measured enzyme activity) are tagged at their corresponding positions. The first and the second principal components (PC1 and PC2) are shown with their contribution rates in parentheses.

To further visualize ML-guided directed evolution in sequence space, we conducted principal component analysis of the variants based on their sequence-derived feature vectors used in the ML model (Figure 6B). In the case of 5M+, ML guided the evolution around 5M, and the highest activity variant was discovered at the position slightly distant from 5M. In contrast, in the case of 5M−, the evolution guided by ML allowed us to find a different region of sequence space with high activity variants around the wild type. These results indicate that ML guided the evolution to the distinct regions of sequence space depending on the presence/absence of 5M in training data.

## Discussion

In this study, ML-guided directed evolution of SrtA was performed to investigate the effects of a highly positive variant 5M in training data. The two series of ML-guided directed evolution with and without 5M explored the distinct regions of sequence space, and both discovered the variants with higher enzyme activity than 5M. These series correspond to typical scenarios encountered in protein engineering: (1) we already have a highly positive variant that has a desired function at least to some degree; and (2) no such variant is available. Our results suggest that ML is useful in both scenarios for discovering improved variants, and each scenario can provide variants with distinct sequences, expanding the sequence diversity of improved variants.

In ML prediction, we ranked all possible five-point mutants of SrtA, thus the size of sequence space was 20^5^ = 3,200,000. In our previous study [8], we performed ML-guided directed evolution of fluorescent proteins to explore sequence space of all four-point mutants (20^4^ = 160,000). In both studies, the initial training data consisted of approximately 100 variants, of which only 3–5% were functionally positive (e.g., the 4 variants with higher enzyme activity than WT in Figure 3). Nonetheless, the libraries proposed by ML were enriched with high performance variants. Therefore, this size of initial training data should be sufficient for discovering high performance variants from the sequence space of 20^4^–20^5^.

In previous applications of ML in mutagenesis-based protein engineering, training data were prepared by two different methods: (1) each variant is isolated, and its function is measured individually [8,11-15]; and (2) functions are measured from a mixture of variants via certain information that may correlate with the functions, e.g., read counts in deep sequencing [9,10,26-28]. The quality of training data obtained by the former method can be high while their size is usually limited to tens or hundreds. Thus, this type of training data is suitable when the number of mutated residues is relatively small, as in our present study. The latter method can provide larger training data with the size of thousands or tens of thousands while their quality may be lower due to the indirect measurement of function such as read counts. In addition, the latter method can only be applied to specific kinds of proteins such as antibodies for which assays from a mixture of variants are established.

The 5M variant was previously discovered by high-throughput screening of SrtA variants using yeast display and fluorescence-activated cell sorting [22]. The library size in this previous study was approximately 10^8^. Here, we discovered novel variants with higher enzyme activity than 5M from the region of sequence space around 5M (Figure 6). This indicates that a highly positive variant discovered by massive experimental screening is not always optimal, and ML can discover further improved variants. Moreover, our result using training data without 5M showed an additional ability of ML to discover improved variants from another region of sequence space distant from 5M. These results suggest multiple executions of ML using subsets of the whole training data as a means to expand the sequence diversity of improved variants.

In conclusion, our study demonstrated the importance of regulating the composition of training data in ML-guided directed evolution. Inclusion of a highly positive variant in training data is not always necessary for discovering improved variants. In addition, multiple executions of ML with subsets of training data may discover distinct regions of sequence space containing improved variants.

## Methods

### Preparation of SrtA variants for screening

To generate the initial library, the five residues (94, 160, 165, 190, and 196) were mutated by the method described previously [8]. Briefly, point saturation mutagenesis at each residue, and site-directed random mutagenesis at all five residues were performed by means of 22-c trick method [29]. The gene fragments of the SrtA variants were generated from the plasmid containing the wild-type sequence by overlap extension PCR [30] using external and 22-c trick primers. The second- and third-round libraries proposed by ML were generated in a similar way using appropriate mix primers instead of 22-c trick primers. The variant gene fragments were ligated into pET22b vectors, and *E. coli* (DE3) bacteria were transformed with the resultant vectors. Colonies of transformed bacteria grown on agar media plates containing 100 µg/mL ampicillin were randomly picked up, incubated overnight at 37 °C in 1 mL of LB broth containing 100 μg/mL ampicillin in a deep-well plate (Axygen, CA, USA), and used for gene sequence analysis. A 100-µL aliquot of each cell culture was added into 900 µL of 2× YT broth supplemented with 100 μg/mL ampicillin in a deep-well plate and further cultured. When the optical density of the culture medium reached 0.6–0.8, isopropyl-1-thio-L-D-galactopyranoside was added to each well to a final concentration of 1 mM to induce SrtA variant expression, and the cells were further cultured for 3 h.

### Screening of SrtA variants in the libraries

The harvested cells in each well were centrifuged, and the cell pellets were lysed in 200 µL of Bugbuster Master Mix (Merk Millipore, Damstadt, Germany). After centrifugation, the expressed SrtA variant in each supernatant was immobilized on an IMAC column, washed with 50 mM imidazole solution, and eluted with buffer A (50 mM Tris-HCl pH 8.0, 200 mM NaCl) containing 300 mM imidazole. Concentrations of the IMAC-purified SrtA variants were estimated by means of the Bradford method. LPETG-fused GFP and GGG-fused YFP were expressed in *E. coli* and purified by means of IMAC and SEC. For the screening assay of enzyme activity, 20 µM of the two substrates (LPETG-fused GFP and GGG-fused YFP) were reacted in buffer A containing 5 mM CaCl_2_ and 3 ng/µL IMAC-purified SrtA (wild type or variant) for 12 h at 37 °C. Then, 0.5 M ethylenediaminetetraacetic acid was added to stop the reaction, and a 10-µL aliquot of each solution was analyzed by SDS-PAGE. The amount of the conjugate (YFP-linked GFP) relative to the control protein was quantified with Image Quant TL software.

To evaluate the enzyme activity of selected SrtA variants in detail, the transformed cells were cultivated in a 1-L shake flask containing 500 mL of 2× YT broth supplemented with 100 μg/mL ampicillin. The expression of the SrtA variant was induced by adding 1 mM isopropyl-1-thio-L-D-galactopyranoside when the optical density of the culture medium reached 0.6, and the cells were grown overnight. The harvested cells were centrifuged, and the pellet was suspended in buffer A and ultrasonicated. The suspension was centrifuged, and the supernatant was purified by means of IMAC and SEC. Then 200 µM GGG-fused YFP was mixed with 0–120 µM LPETG-fused GFP in buffer A containing 5 mM CaCl_2_ and 3 ng/µL purified SrtA, and the enzyme activity was measured as described for the SrtA screening assay above.

### ML model

We previously used the Bayesian optimization software COMBO [31] to conduct ML-guided directed evolution of fluorescent proteins [8]. Here, we applied a similar procedure for enhancing the enzyme activity of SrtA. COMBO implements a Gaussian process based on a linear regression model using a random feature map:

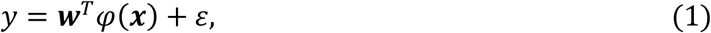

where *y* is the enzyme performance score of the protein (defined in the next section), ***x*** is a feature vector of the protein, *φ*(***x***) is a random feature map, and *ε* is an error term. Given a training dataset {(*y*, ***x***)}, COMBO fits a weight vector ***w*** so that the enzyme performance score *y* can be predicted from the feature vector ***x***. For each variant not included in the training dataset, COMBO computes the acquisition function by Thompson sampling [31], which evaluates the probability that the enzyme performance score of the variant is higher than any variant in the training dataset. These values were used to rank all possible variants in sequence space.

Here, we performed two rounds of ML for each of two scenarios (5M+ and 5M−). In the first round, two different training datasets were prepared (initial 5M+ and 5M− libraries in Table 1; Supplementary Table S1), and separately used to construct two ML models. These ML models were used for ranking all possible variants, obtaining the two ranking lists (Supplementary Data S1 for 5M+; Supplementary Data S2 for 5M−). In the second round, the top-ranked variants from the first-round ML were used as additional training data (Supplementary Table S2) to update the ML models and the ranking lists (Supplementary Data S3 for 5M+; Supplementary Data S4 for 5M−).

### Enzyme performance score

To discover improved variants, the ML model needs to predict variants that have not only high enzyme activity but also expression levels sufficient for measuring enzyme activity. To satisfy this requirement, the following enzyme performance score was used in the ML model:

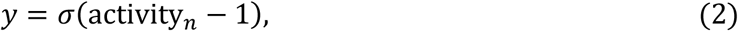

where *σ*(·) is a sigmoidal function, and activity_*n*_ is the enzyme activity of the protein divided by that of the wild type. Importantly, we set activity_*n*_ as zero if the expression level is insufficient for measuring enzyme activity. Thus, the enzyme performance score takes a high value when the enzyme activity is high while it takes the minimum value for variants with insufficient expression levels. This allows the ML model to predict high activity variants while avoiding variants with insufficient expression levels.

### Feature vector

We defined a feature vector ***x*** of the SrtA protein by concatenating the feature vectors of amino acids at the five mutated residues. For a feature vector of each amino acid, we considered a variety of amino acid descriptors including Z-scale [24], T-scale [32], ST-scale [33], FASGAI [34], MS-WHIM [35], ProtFP [36], VHSE [37], and BLOSUM-based features [38]. In addition, we used PSSM as a feature vector to incorporate the evolutionary information from SrtA homologs. Specifically, PSSM was constructed by homology search using PSI-BLAST [25] with the wild-type SrtA sequence as a query against the nr database. The homology search was iterated five times with 200 homologs used for updating PSSM per iteration. This produced PSSM {*s*_*ij*_} for each residue *i* and amino acid *j* (Supplementary Data S5). We constituted a PSSM-based feature vector of the SrtA protein by concatenating the PSSM values at the five mutated residues and their amino acids.

To select the optimal feature vector, we performed a feature selection procedure like that in our previous study [8]. Briefly, we performed a benchmark experiment where COMBO was set to find 5M from the initial library by a Bayesian optimization procedure (Supplementary Figure S4). In this experiment, MS-WHIM, Z-scale, and PSSM achieved better results than the other descriptors in terms of the number of training data needed to find 5M. These three descriptors were further compared in another benchmark experiment where COMBO was trained using the initial library with 5M excluded (i.e., initial 5M− library), and all possible variants in sequence space of 20^5^ were ranked. In this experiment, the combination of Z-scale and PSSM achieved the best result in terms of the rank of 5M (Supplementary Table S4). Therefore, we used the combination of Z-scale and PSSM as the feature vector for our final model in all other parts of this study.

## Acknowledgements

We thank Ms. Miho Hosoya, Ms. Hiromi Ogata, and Ms. Yuri Ishigaki for experimental support. This work was partly supported by the Cross-ministerial Strategic Innovation Promotion Program (SIP) “Technologies for Smart Bio-industry and Agriculture” (funding agency: Bio-oriented Technology Research Advancement Institution, National Agriculture and Food Research Organization (NARO), Japan); Scientific Research Grant from the Japan Society for the Promotion of Science Research Fellowships (JP16H04570, JP16K14483, JP20H00315 to M.U.); and the project “Development of the Key Technologies for the Next-Generation Artificial Intelligence/Robots” of the Ministry of Economy, Trade and Industry, Japan (to M.U.). Computations were partly performed on the NIG supercomputer at ROIS National Institute of Genetics, AI Bridging Cloud Infrastructure (ABCI) at National Institute of Advanced Industrial Science and Technology (AIST), and the supercomputer of ACCMS at Kyoto University.

## Author Contributions

YS conducted the computational analysis. MO, TS, HK, and TI conducted the experiments. YS, MO, TK, KT and MU participated in the data interpretation. YS and MU wrote the paper. KT and MU conceived of the study and directed the project. All authors read and approved the final version of the manuscript.

## Supplementary Material

Supplementary Information contains Supplementary Tables S1-S4, Supplementary Figures S1-S4, and the legends for Supplementary Data S1-S5. The bodies of Supplementary Data S1-S5 are available as separate Excel files.

