## Supplementary Information for "Machine-learning-guided library design cycle for directed evolution of enzymes: the effects of training data composition on sequence space exploration"

### Supplementary Tables and Figures

**Supplementary Table S1.** SrtA variants in the initial library. Sequence information is shown by five amino acids at the mutated residues (94, 160, 165, 190, and 196). Measured: variants with measured enzyme activity. Low exp.: variants with insufficient expression levels for measuring enzyme activity.

| | |
|---|---|
| Wild type | PDDKK |
| 5M | RNAET |

Variants

| Random mutagenesis | | Point mutagenesis | | | |
|---|---|---|---|---|---|
| Measured | Low exp. | Measured | | Low exp. | |
| HDFDK | LTDFQ | 94D | DDDKK | 160V | PVDKK |
| QGDAK | SKAWK | 94F | FDDKK | 195P | PDDKP |
| EQLGM | GDYAV | 94I | IDDKK | | |
| CYGPE | IYDVM | 94L | LDDKK | | |
| KGFVV | MDDYI | 94M | MDDKK | | |
| TDWSV | NGDND | 94T | TDDKK | | |
| TWDNL | AVNYL | 94V | VDDKK | | |
| NGDDD | HHRVM | 160F | PFDKK | | |
| HGFNK | PLDCC | 160K | PKDKK | | |
| YRDKG | HADEF | 160T | PTDKK | | |
| LPDNR | MLDVF | 165H | PDHKK | | |
| LGDDG | ACDWE | 165K | PDKKK | | |
| VKDPQ | ELDIQ | 165L | PDLKK | | |
| NYDVH | CYGVE | 190I | PDDIK | | |
| VSDGI | NDKYI | 190L | PDDLK | | |
| WEDNL | KPKEF | 190P | PDDPK | | |
| WDDAL | EFDVA | 190V | PDDVK | | |
| SIDWQ | VYDEI | 190Y | PDDYK | | |
| DYKHH | QCDWL | 195C | PDDKC | | |
| MWDMG | FWDFK | 195Q | PDDKQ | | |
| TWDQG | IADSL | 195R | PDDKR | | |
| HQTVK | YVDFQ | 195V | PDDKV | | |
| HQTDK | DNDWV | | | | |
| | TTIPY | | | | |
| | FWSGG | | | | |
| | SWDLG | | | | |
| | TWDWP | | | | |
| | ACYEP | | | | |
| | CDYTG | | | | |
| | IGDWT | | | | |
| | TAQNR | | | | |
| | AFDLR | | | | |
| | YVDLL | | | | |

**Supplementary Table S2.** SrtA variants in the second-round libraries. Sequence information is shown by five amino acids at the mutated residues (94, 160, 165, 190, and 196). Also shown are ranks by ML prediction. Ranks without sequences (—) mean that these variants could not be cloned from the library. Sequences in parentheses represent variants with insufficient expression levels for measuring enzyme activity.

Second-round 5M+ library

| Top 50 | | | | Lower than top 50 | |
|---|---|---|---|---|---|
| Rank | Sequence | Rank | Sequence | Rank | Sequence |
| 1 | RNAQT | 31 | RNPKT | 51 | KNAKT |
| 2 | PRDKK | 32 | RDAKT | 52 | RDDKT |
| 3 | RNPET | 33 | — | 54 | KNNKT |
| 4 | RNSQT | 34 | PHDKK | 58 | RNPNT |
| 5 | RNPQT | 35 | RDDKK | 64 | RSSET |
| 6 | RNSET | 36 | — | 66 | RDPKT |
| 7 | RDAQT | 37 | RDNET | 67 | PHDRK |
| 8 | RDAET | 38 | KNDKK | 70 | KNPKT |
| 9 | — | 39 | KNSKT | 71 | RNDKK |
| 10 | PRDRK | 40 | RSPQT | 82 | KNDKT |
| 11 | — | 41 | KNNQT | 83 | KDAKT |
| 12 | — | 42 | KDNQT | 86 | RNPEE |
| 13 | RNNQT | 43 | KNSET | 99 | KNNET |
| 14 | KNAET | 44 | — | 100 | KDPET |
| 15 | RNAKT | 45 | — | 105 | RNPEG |
| 16 | RDNQT | 46 | RNAST | 106 | RNDKT |
| 17 | RNSKT | 47 | — | 112 | RDAEG |
| 18 | — | 48 | — | 116 | RDAST |
| 19 | RDPQT | 49 | KDDKT | 117 | RSAKT |
| 20 | — | 50 | KDAET | 118 | RDANT |
| 21 | RSAET | | | 134 | RSPNT |
| 22 | RDPET | | | 140 | RSPKT |
| 23 | — | | | 151 | KDPKT |
| 24 | KDDKK | | | 155 | RDAEE |
| 25 | RNNET | | | 202 | RDPNT |
| 26 | — | | | 215 | RDPST |
| 27 | RSPET | | | 257 | RDPEG |
| 28 | RSAQT | | | 315 | RDPQE |
| 29 | KNPET | | | 554 | RSPTT |
| 30 | KDAQT | | | 606 | RNPKG |

**Supplementary Table S2 (continued).**

Second-round 5M− library

| Top 50 | | | | Lower than top 50 | | | |
|---|---|---|---|---|---|---|---|
| Rank | Sequence | Rank | Sequence | Rank | Sequence | Rank | Sequence |
| 1 | PRDKK | 31 | PKDQK | 52 | PRDQT | 4238 | PKKRK |
| 2 | PRDRK | 32 | PHNKK | 53 | RDDRK | | |
| 3 | PHDKK | 33 | PKDKV | 54 | PKDRV | | |
| 4 | PHDRK | 34 | PKDKR | 57 | PKERK | | |
| 5 | — | 35 | — | 60 | PHDHK | | |
| 6 | PRDQK | 36 | KEDKK | 62 | KEDRK | | |
| 7 | PKDRK | 37 | PKDRT | 65 | PKNRK | | |
| 8 | PREKK | 38 | PHDRR | 67 | PRERR | | |
| 9 | — | 39 | KNDRK | 69 | PHDRQ | | |
| 10 | KDDKK | 40 | PKEKK | 71 | RNDKK | | |
| 11 | PRNKK | 41 | PRDRV | 72 | PRDQQ | | |
| 12 | PRDRR | 42 | PHERK | 73 | PKDKQ | | |
| 13 | — | 43 | PHDRV | 74 | PHSKK | | |
| 14 | PRDKQ | 44 | PREKR | 88 | PHDQT | | |
| 15 | PRDRT | 45 | PKNKK | 93 | PKDRQ | | |
| 16 | PHDKT | 46 | PREQK | 108 | PKSKK | | |
| 17 | KDDRK | 47 | PHDKQ | 110 | PHSRK | | |
| 18 | PRNRK | 48 | PRSRK | 121 | PKDQT | | |
| 19 | — | 49 | PKDRR | 127 | RNDRK | | |
| 20 | PRDHK | 50 | PHNRK | 129 | PHEQK | | |
| 21 | KNDKK | | | 152 | KKDKK | | |
| 22 | PRDRQ | | | 155 | PKDQV | | |
| 23 | PHDKR | | | 157 | PHEKR | | |
| 24 | PRDKV | | | 170 | PKDQR | | |
| 25 | PKDKT | | | 182 | PREQR | | |
| 26 | PHDKV | | | 208 | PHDQQ | | |
| 27 | PHEKK | | | 214 | PHERR | | |
| 28 | RDDKK | | | 237 | KKDRK | | |
| 29 | (PHDRT) | | | 326 | PKDQQ | | |
| 30 | PRSKK | | | 3581 | PKKKK | | |

**Supplementary Table S3.** SrtA Variants in the third-round libraries. Sequence information is shown by five amino acids at the mutated residues (94, 160, 165, 190, and 196). Also shown are ranks by ML prediction. Sequences in parentheses represent variants with insufficient expression levels for measuring enzyme activity.

| Third-round 5M+ library | | | | Third-round 5M− library | | | |
|---|---|---|---|---|---|---|---|
| Rank | Sequence | Rank | Sequence | Rank | Sequence | Rank | Sequence |
| 1 | KSSKT | 31 | KNQQT | 1 | (PKEQR) | 31 | PRTRK |
| 2 | KSAKT | 32 | KSNET | 2 | PHEQR | 32 | PRDRL |
| 3 | RSSKT | 33 | KSAKE | 3 | PQEQR | 33 | PQDRR |
| 4 | KNAQT | 34 | KSSKE | 4 | PKEKR | 34 | PHDKE |
| 5 | KAAKT | 35 | QNAKT | 5 | PKNQR | 35 | PMEQR |
| 6 | KSAQT | 36 | RNEKT | 6 | PQEKR | 36 | PHARK |
| 7 | KASKT | 37 | RGSKT | 7 | PQEKK | 37 | PTEQR |
| 8 | RSQKT | 38 | RSGKT | 8 | PKQQR | 38 | PKEQK |
| 9 | KSTKT | 39 | QSSKT | 9 | RKDKK | 39 | KRDKK |
| 10 | QSAKT | 40 | KSDKT | 10 | PKERR | 40 | PQNQR |
| 11 | KQSKT | 41 | KSQQT | 11 | PRDQV | 41 | PHSQR |
| 12 | KSSRT | 42 | KSEKT | 12 | PKEER | 42 | PRKRK |
| 13 | KSGKT | 43 | RTSKT | 13 | (PHQQR) | 43 | KNDRR |
| 14 | KSQKT | 44 | KGAKT | 14 | PKQRK | 44 | PQNQK |
| 15 | RNNKT | 45 | RSSKE | 15 | PKNKR | 45 | PHEER |
| 16 | RSEKT | 46 | KQAQT | 16 | PQNKK | 46 | PQERR |
| 17 | (KSSQT) | 47 | RNANT | 17 | PRDRI | 47 | PKRRK |
| 18 | KSSKQ | 48 | RNADT | 18 | QNDRK | 48 | PKKRR |
| 19 | KQAKT | 49 | RSDKT | 19 | RQDKK | 49 | PQDKK |
| 20 | RSTKT | 50 | KTAKT | 20 | TNDRK | 50 | PQDQR |
| 21 | RSNKT | | | 21 | PQDKR | | |
| 22 | KSAKQ | | | 22 | PRERK | | |
| 23 | RASKT | | | 23 | PQNKR | | |
| 24 | RSSKQ | | | 24 | PQEQK | | |
| 25 | KSART | | | 25 | PKNQK | | |
| 26 | KSNKT | | | 26 | PHDRI | | |
| 27 | RSAST | | | 27 | PRDQI | | |
| 28 | (RSSRT) | | | 28 | PKDQI | | |
| 29 | KNSQT | | | 29 | PKQRR | | |
| 30 | RESKT | | | 30 | PHDRL | | |

**Supplementary Table S4.** Benchmark of feature vectors. For each type of feature vector, the ML model was trained using the initial 5M− library, and used to rank 5M among all unknown variants in the sequence space of $20^5$ = 3,200,000. M+P: feature vectors of MS-WHIM and PSSM are concatenated. Z+P: feature vectors of Z-scale and PSSM are concatenated.

| Feature vector | Rank of 5M |
|---|---:|
| MS-WHIM | 704,598 |
| Z-scale | 445,624 |
| PSSM | 1,957,838 |
| M+P | 472,521 |
| Z+P | 161,052 |

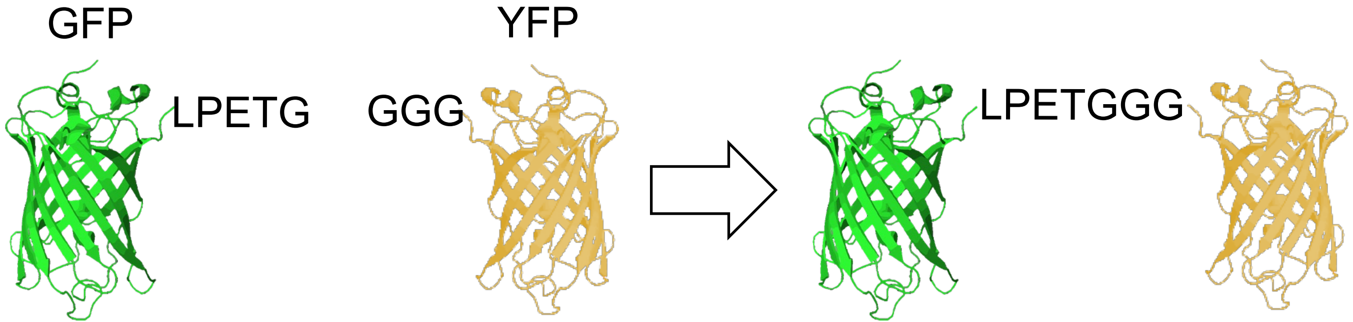


**Supplementary Figure S1.** Cross-linking reaction of GFP and YFP catalyzed by SrtA. GFP with an LPETG sequence at the C-terminus and YFP with a GGG sequence at the N-terminus were conjugated with each other by a SrtA variant.

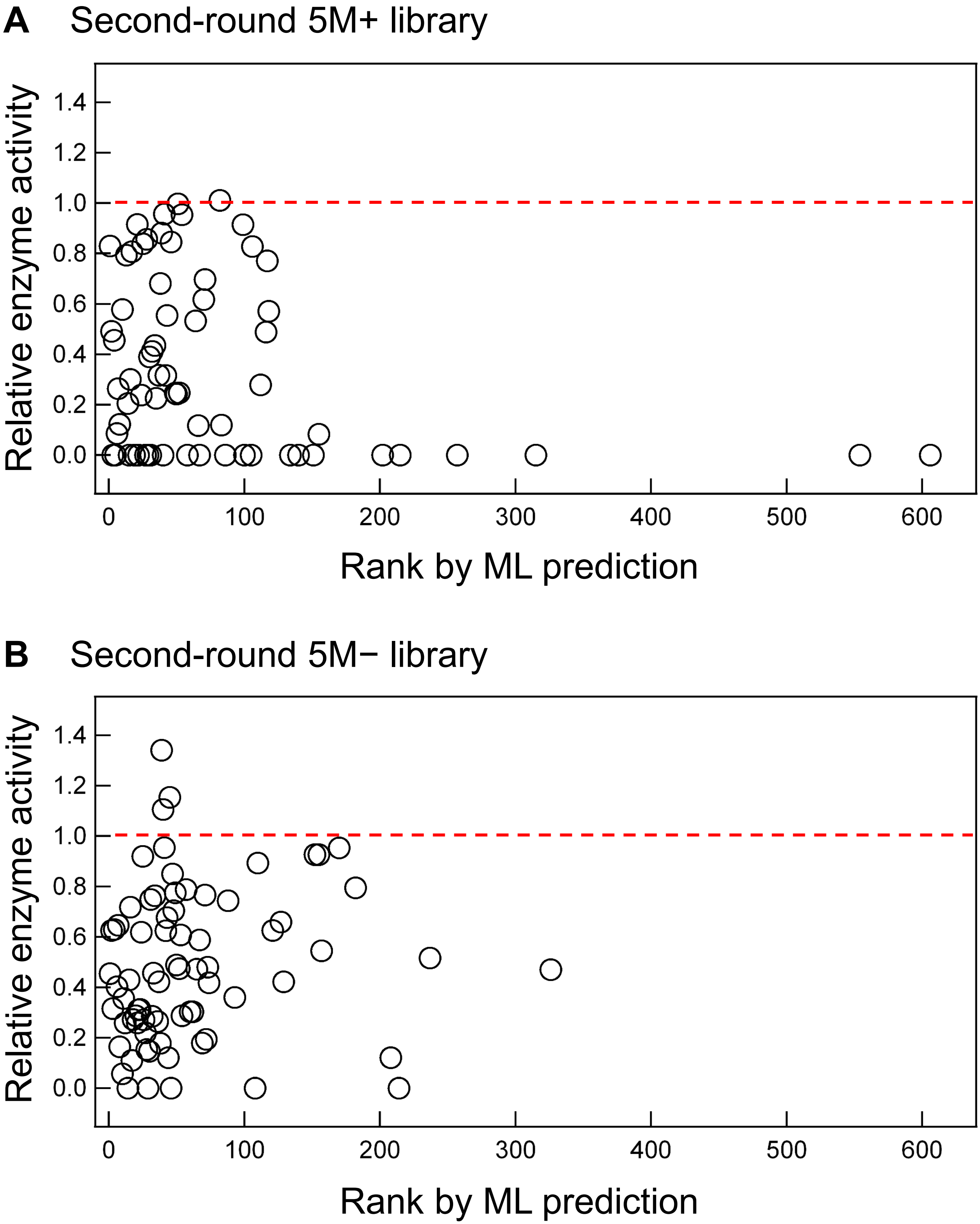


**Supplementary Figure S2.** Enzyme activity of the SrtA variants in the second-round libraries versus their ranks by ML prediction. Enzyme activity is normalized by that of 5M (red dashed line).

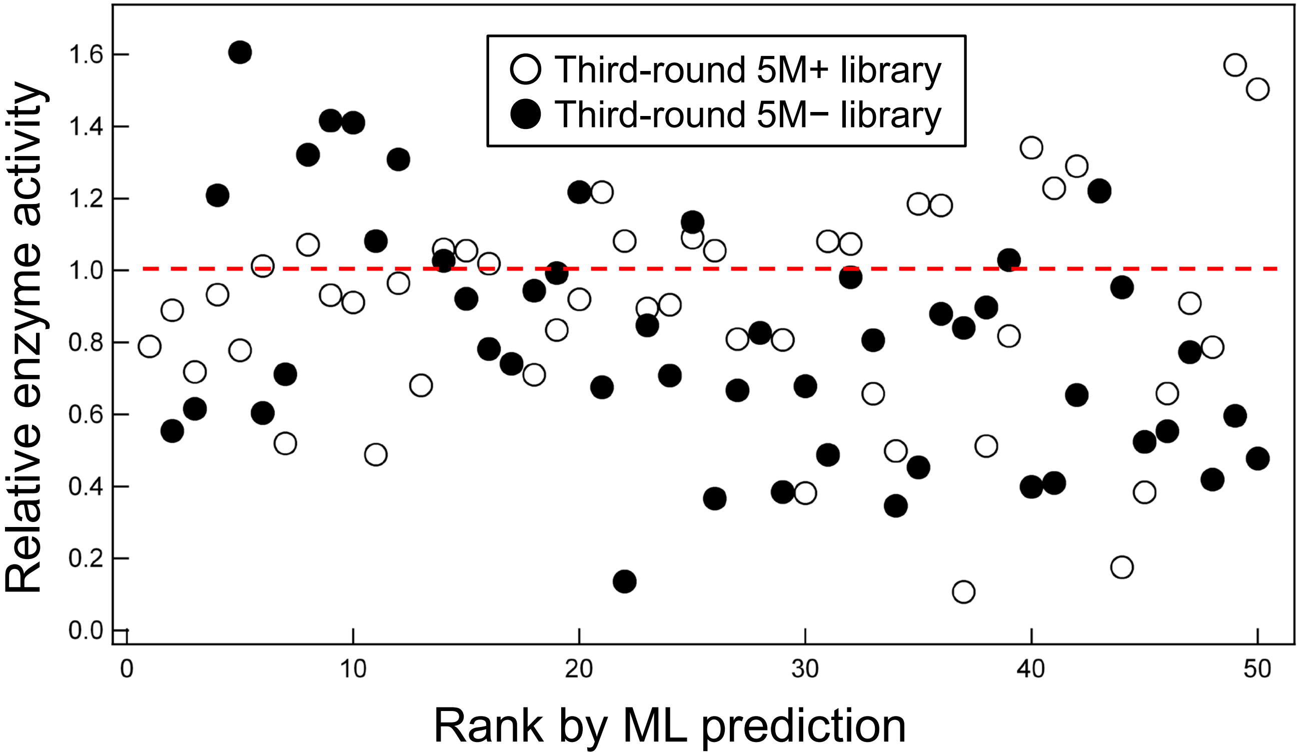


**Supplementary Figure S3.** Enzyme activity of the SrtA variants in the third-round libraries versus their ranks by ML prediction. Enzyme activity is normalized by that of 5M (red dashed line).

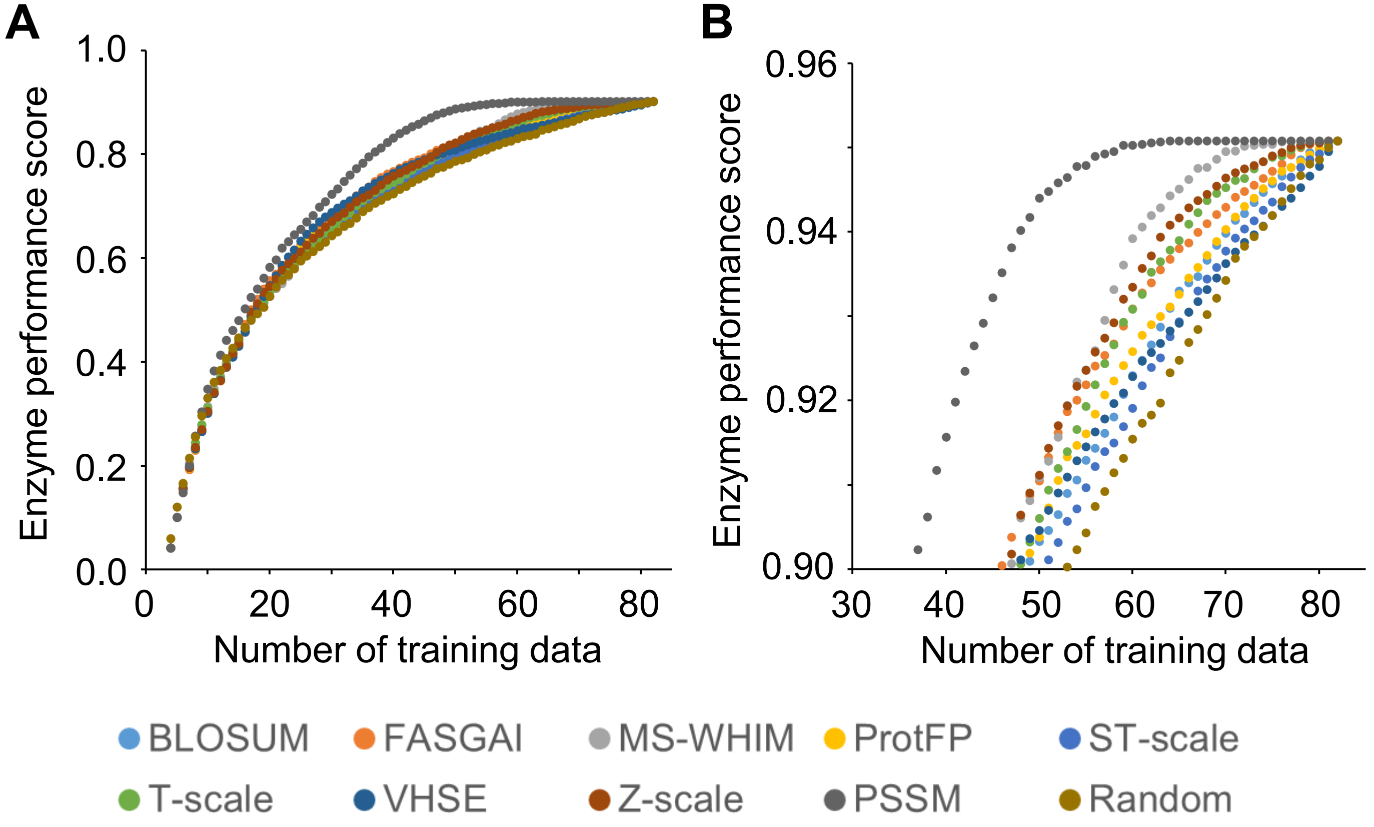


**Supplementary Figure S4.** Benchmark of feature vectors. (A) For each type of feature vector, Bayesian optimization was conducted to find variants with high enzyme performance scores among the initial 5M+ library. The enzyme performance score of the best variant found is plotted against an increasing number of training data. The Bayesian optimization procedures were repeated 1,000 times with different choices of initial training data points (N=5), and the average results are shown. Feature vectors achieving higher enzyme performance scores at smaller numbers of training data are regarded as good feature vectors. Z-scale, MS-WHIM, and PSSM showed better performance than the other feature vectors. (B) Magnified version of (A).

### Supplementary Data

**Supplementary Data S1.** Ranking list by ML prediction using the initial 5M+ library as training data. This list was used for designing the second-round 5M+ library. The top 160,000 variants are shown. The data are available as a separate Excel file.

**Supplementary Data S2.** Ranking list by ML prediction using the initial 5M− library as training data. This list was used for designing the second-round 5M− library. The top 160,000 variants are shown. The data are provided as a separate Excel file.

**Supplementary Data S3.** Ranking list by ML prediction using the second-round 5M+ library as additional training data. This list was used for designing the third-round 5M+ library. The top 160,000 variants are shown. The data are provided as a separate Excel file.

**Supplementary Data S4.** Ranking list by ML prediction using the second-round 5M− library as additional training data. This list was used for designing the third-round 5M− library. The top 160,000 variants are shown. The data are provided as a separate Excel file.

**Supplementary Data S5.** PSSM of SrtA homologs. This was used to calculate PSSM-based feature vectors for ML. The data are provided as a separate Excel file.
